# Coring has no detrimental impact on tree growth across European forests, but effects on mortality remain uncertain

**DOI:** 10.64898/2026.02.03.703454

**Authors:** Robin Battison, Thomas S. Ovenden, Daniela Nemetschek, Fabian Jörg Fischer, Olivier Bouriard, Giovanni Iacopetti, Tommaso Jucker

**Affiliations:** School of Biological Sciences, University of Bristol, Bristol, BS8 1TQ, UK; Forest Research, Northern Research Station, Roslin, Midlothian EH25 9SY, UK; Technical University of Munich, School of Life Sciences, Ecosystem Dynamics and Forest Management, Hans-Carl-von-Carlowitz-Platz 2, 85354 Freising, Germany; Ștefan cel Mare University of Suceava, Str. Universității 13, 720229 Suceava, Romania; Università di Firenze, Piazza di San Marco, 4, 50121 Firenze FI, Italy

**Author notes:** **Corresponding authors**: Robin Battison and Tommaso Jucker.

**Keywords:** Dendrochronology, forest census data, tree coring, tree growth, tree mortality, coring impact

## Abstract

1. Tree cores are widely used across a broad range of disciplines in the environmental sciences, most notably as a tool to measure tree growth, estimate tree age, characterise wood anatomy and reconstruct past climate. However, because extracting tree cores is an invasive procedure, concerns about their use are often raised due to perceived risks to trees.
2. Here we comprehensively test the long-term impacts of tree coring on 16 European tree species using a dataset spanning a broad range of forest types across the continent. Over the course of a decade, we tracked the growth and survival of 3201 trees cored in 2012 (including those cored once and twice) and used hierarchical Bayesian models to compare them with those of a cohort of 7262 neighbouring trees that were never cored.
3. We found no evidence that coring had a detrimental impact on tree growth, irrespective of size, species, forest type or the number of times a tree was cored. However, we did observe a small positive stem increment response, which we hypothesise is most likely the result of localised vertical scarring from the coring wound.
4. Impacts of tree coring on mortality were instead more uncertain, with mortality rates of cored trees slightly elevated relative to ones that were not cored, but overlapping widely in their credible intervals. This underscores the challenge of attributing drivers of rare and stochastic processes such as tree mortality, even with relatively large datasets.
5. Our study generally supports the use of tree coring as a low-impact method for characterising the growth, age and function of a wide range of tree species, but highlights that there are circumstances in which it may cause harm. Future work should explore these further by considering the broader impacts of coring on tree health. Additionally, tree cores should always be collected well above or below the point of measurement of tree stem diameters to avoid biasing long-term forest census datasets.

## Introduction

Tree coring is central to the science of dendrochronology and is used widely in ecology as a tool to reconstruct the age, growth rate and the ecophysiological status of trees. Data extracted from tree cores are used to characterise how tree growth rates vary through time and in response to biotic and abiotic drivers (Anderson-Teixeira et al., 2022; Jucker, Bouriaud, Avacaritei, Dănilă, et al., 2014) and quantify forest productivity and carbon storage (Babst et al., 2014; Li et al., 2024). Tree cores are also useful to understand how trees respond to climate extremes (Anderegg et al., 2015; Castagneri et al., 2022; Sohn et al., 2016; Wei et al., 2024), measure wood traits related to plant hydraulics and physiology (Fonti et al., 2009; Grossiord et al., 2014) and reconstruct past climate (Emile-Geay et al., 2017; Esper et al., 2016; Rodriguez-Caton et al., 2024). Moreover, historical growth reconstruction from tree rings is increasingly being combined with other spatio-temporal datasets, such as satellite imagery, to track forest responses to climate change at scale (Kannenberg et al., 2019; Levesque et al., 2019; Mašek et al., 2024; Mathes et al., 2024; Morin-Bernard et al., 2024). However, due to its invasive nature, there have long been concerns about the potential adverse effects it could have on trees (Tsen et al., 2016) and because of this, a precautionary approach has meant tree coring is often not permitted in conservation areas and in long-term ecological monitoring sites.

Tree cores are extracted using an increment borer, which leaves behind a hole in the trunk that is typically 5–10 mm in diameter (depending on the increment borer used) and extends from the outer bark to the centre of the tree. These bore holes are thought to potentially impact tree growth, mortality risk, and physiology in several ways. First and foremost, they can serve as a site of infection, particularly for fungal pathogens in warm, wet climates (Boura et al., 2014; Florens, 2013, 2014). These infections can lead to wood staining (most likely from the release of tannins; (Kelsey & Harmon, 1989) and can result in wood decay, although from the limited evidence available this appears to be more common in broadleaf trees (e.g., *Betula* and *Populus* species) with little evidence of it occurring in conifers (Tsen et al., 2016; Wunder et al., 2013). To prevent infection it is relatively common practice to plug wounds with synthetic materials or antifungal compounds, although there is little evidence that this has an effect and some concern it may increase risk of infection if it traps moisture (Dujesiefken et al., 1999). The wound caused by extracting the cores can also lead to a localised growth response by trees. For example, *Picea abies* was shown to have elevated localised growth near coring sites due to vertical scarring and callous tissue overgrowing the wound (Fabiánová & Šilhán, 2021), and large, localised scarring responses have also been reported in *Fagus sylvatica* (Portier et al., 2023). The effects of tree coring on tree vitality may be particularly pronounced in smaller trees, for which bore holes represent a proportionally much larger wound and which may have fewer resources to repair damaged tissue (Datta et al., 2025). Collectively, these studies highlight that further evidence is needed to determine the impacts of coring on tree growth, survival and wood quality (Holl & Road, 2018; Tsen et al., 2016).

Our current understanding of the long-term effects of coring on tree growth and mortality is pieced together from a relatively small number of studies that have focused on a limited number of tree species and study sites (Helcoski et al., 2018; Mantgem & Stephenson, 2004; Portier et al., 2023; Wunder et al., 2011). In particular, only a handful of studies have explored the effects of coring on tree mortality. These studies leveraged large sample sizes (>500 cored trees) and found no adverse effects over a 7–40 year period (Helcoski et al., 2018; Mantgem & Stephenson, 2004; Wunder et al., 2011), but were all conducted at a single location, and only one explored the effects of coring across a significant number (19) of species (Helcoski et al., 2018). Similarly, to the best of our knowledge only one study has investigated the long-term impacts of coring on tree growth, reporting no decrease in growth in the 10 years after coring in three common European tree species (*Picea abies*, *Abies alba*, and *Fagus sylvatica*) sampled in Switzerland and the Ukraine (Portier et al., 2023). This limited body of evidence suggests that coring is unlikely to substantially impact trees. However, we currently lack a comprehensive and systematic assessment of the long-term impacts of coring on tree growth and mortality that not only captures responses across a diverse range of species but also tests whether effects differ among climatically distinct forest types, tree sizes, and coring practices (e.g., extracting two cores per tree to improve cross dating).

Here, we provide the first comprehensive overview of the long-term impacts of tree coring on the growth and survival of 16 common European tree species, comprising 12 broadleaves and four conifers. Over the course of a decade, we tracked the growth and survival of 10,463 trees – 3201 of which were cored in 2012 – distributed across 205 permanent forest plots covering Europe’s major forest types. This includes trees that were cored once (2416) or twice (785) spanning a broad range of sizes (7.5–101.5 cm in stem diameter). Using this unique dataset, we compared the growth and survival of cored and non-cored trees from the same plots and set out to test the following hypotheses:

1. Based on previous studies (Helcoski et al., 2018; Mantgem & Stephenson, 2004; Portier et al., 2023; Wunder et al., 2011), we do not expect tree coring will have a detrimental effect on the long-term growth and survival of most European tree species and forest types.
2. If a detrimental effect of coring is observed, we expect this negative impact will be disproportionately pronounced in smaller trees, as these are more vulnerable to internal decay and wounds (Datta et al., 2025).
3. Whilst often cautioned against (Portier et al., 2023; Tsen et al., 2016), we do not expect that collecting a second core will result in any additional negative impact on tree growth or survival compared to taking a single core, as the size of the wounds remains very small relative to that of tree.

## Methods

### FunDivEUROPE forest plot network

Our analyses are based on data from the FunDivEUROPE permanent forest plot network (Baeten et al., 2013). The network consists of 209 plots (30×30 m in size) distributed across six sites spanning a broad range of biomes and forest types, including Mediterranean, temperate and boreal forests (Fig. 1). The plots cover a latitudinal gradient of more than 20°, ranging from Mediterranean forests in Spain (mean annual temperature and rainfall = 10.2 **°**C and 499 mm) and Italy (13.0 **°**C and 850 mm), temperate forests in Germany (6.8 **°**C and 775 mm) and Romania (6.8 **°**C and 800 mm), hemiboreal forests in Poland (6.9 **°**C and 627 mm) and boreal forests in Finland (2.1 **°**C and 700 mm) (Baeten et al., 2013). The network was established in 2012 within mature forest stands with no recent management history prior to plot establishment and has since been re-censused in 2017 and 2022. Of the original 209 plots, 205 were surveyed across all three census periods, with the remaining four plots abandoned due to harvesting activities that occurred after 2012.

**Fig. 1:**
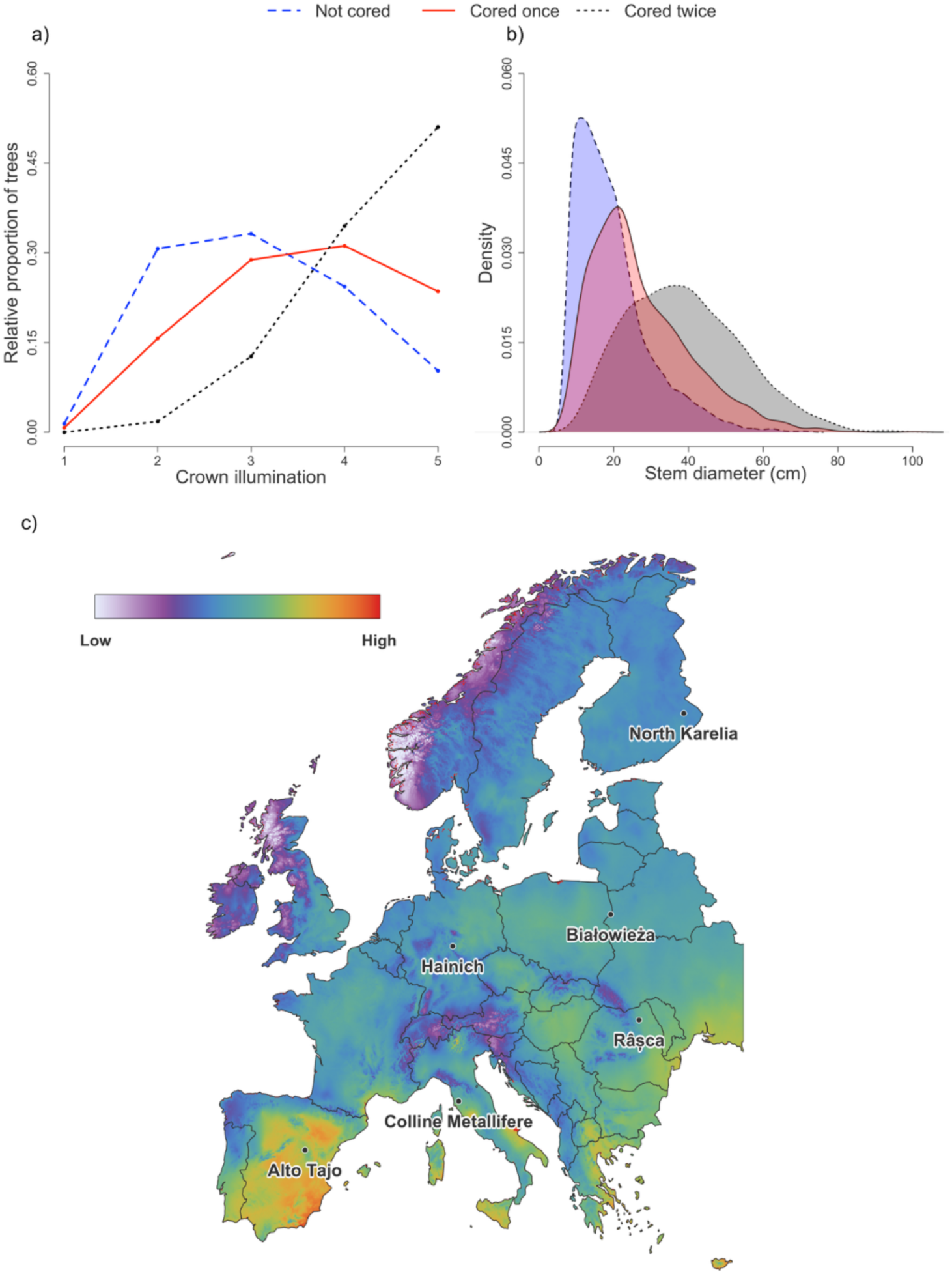
Variation in the distribution of (**a**) crown illumination index scores and (**b**) stem diameters of trees that were either not cored (blue, dashed line), cored once (red, continuous line), or cored twice (black, dotted line). The map (**c**) shows the locations of the FunDivEUROPE sites, with the underlying colour gradient reflecting the aridity index (Zomer et al., 2022).

Within each plot and at each census interval, all stems with a diameter at breast height (DBH, measured in cm at 1.3 m aboveground) greater than 7.5 cm were tagged, measured with a diameter tape or callipers, identified to species and their survival was recorded. In addition, the crown illumination index (CII) of each tree was recorded as a measure of light availability to the crown that varies from 1 (completely suppressed) to 5 (fully exposed) (Clark & Clark, 1992). In total, we tracked the growth and survival of 10,463 trees belonging to 16 common European tree species, several of which were sampled at more than one site (Table 1). This includes four conifers (*Abies alba*, *Picea abies*, *Pinus nigra*, *Pinus sylvestris*) and 12 broadleaves (*Acer pseudoplatanus*, *Betula pendula*, *Carpinus betulus*, *Castanea sativa*, *Fraxinus excelsior*, *Fagus sylvatica*, *Ostrya carpinifolia*, *Quercus cerris*, *Quercus faginea*, *Quercus ilex*, *Quercus petraea*, *Quercus robur*).

**Table 1:** Overview of the data analysed in this study, including the number of plots and trees cored for each species across sites, their survival rates between 2012 and 2022 and summary statistics about their mean growth rates and size.

| Country | Species | Cored |  | Not Cored |  | Mean basal area growth (cm<sup>2</sup> year<sup>-1</sup>) | Mean DBH in 2012 (cm) |
| --- | --- | --- | --- | --- | --- | --- | --- |
|  |  | Total number | Dead (%) | Total number | Dead (%) |  |  |
| Finland<br>n plots = 28 | *Picea abies* | 203 | 5.4 | 654 | 4.6 | 10.3 | 17.8 |
|  | *Pinus sylvestris* | 176 | 15.3 | 473 | 18.6 | 10.2 | 19.3 |
| Poland<br>n plots = 41 | *Betula pendula* | 77 | 35.1 | 112 | 22.3 | 19.4 | 36.0 |
|  | *Carpinus betulus* | 132 | 6.1 | 717 | 7.1 | 8.2 | 18.8 |
|  | *Picea abies* | 119 | 79.0 | 249 | 63.1 | 11.5 | 28.0 |
|  | *Pinus sylvestris* | 101 | 9.9 | 130 | 6.2 | 28.4 | 41.3 |
|  | *Quercus robur* | 105 | 5.7 | 101 | 5.0 | 31.2 | 35.3 |
| Germany<br>n plots = 38 | *Acer psudeoplatanus* | 83 | 9.6 | 124 | 7.3 | 9.3 | 24.1 |
|  | *Fagus sylvatica* | 238 | 8.4 | 329 | 10.5 | 12.1 | 22.9 |
|  | *Fraxinus excelsior* | 144 | 25.0 | 239 | 36.4 | 15.6 | 26.3 |
|  | *Picea abies* | 67 | 67.2 | 208 | 66.8 | 17.0 | 24.1 |
|  | *Quercus petraea* | 73 | 9.6 | 24 | 8.3 | 26.8 | 50.0 |
| Romania<br>n plots = 28 | *Abies alba* | 95 | 9.5 | 206 | 10.2 | 21.2 | 34.0 |
|  | *Acer psuedoplatanus* | 81 | 12.3 | 152 | 12.5 | 11.4 | 32.0 |
|  | *Fagus sylvatica* | 120 | 13.3 | 336 | 11.6 | 11.3 | 28.0 |
|  | *Picea abies* | 87 | 12.6 | 168 | 16.1 | 19.0 | 36.4 |
| Italy<br>n plots = 36 | *Castanea sativa* | 144 | 52.8 | 410 | 52.0 | 4.2 | 18.0 |
|  | *Ostrya carpinifolia* | 132 | 16.7 | 469 | 23.0 | 3.6 | 14.0 |
|  | *Quercus cerris* | 161 | 9.3 | 244 | 23.0 | 9.8 | 22.9 |
|  | *Quercus ilex* | 166 | 1.8 | 262 | 6.5 | 5.4 | 19.3 |
|  | *Quercus petraea* | 142 | 11.3 | 178 | 23 | 9.7 | 22.6 |
| Spain<br>n plots = 34 | *Pinus nigra* | 165 | 0.6 | 336 | 1.5 | 4.8 | 22.5 |
|  | *Pinus sylvestris* | 104 | 6.7 | 235 | 7.2 | 5.0 | 24.4 |
|  | *Quercus faginea* | 204 | 4.9 | 916 | 4.5 | 2.1 | 12.6 |

### Tree coring

In 2012, when the FunDivEUROPE plots were first established, we acquired tree cores from a subset of trees in each plot to measure their historical growth rates and capture their physiological responses to drought using carbon isotopes (Grossiord et al., 2014; Jucker, Bouriaud, Avacaritei, Dănilă, et al., 2014). In total, 2730 trees were cored for growth reconstruction (approximately 20–25% of trees per plot), while a further 1408 tree cores were collected for isotope analysis (Grossiord et al., 2014; Jucker et al., 2017). Most of the cores collected for isotope analysis overlapped with those sampled for growth, meaning that 805 trees were cored twice. In each case, cores were collected using 5.15 mm diameter increment borers (Haglöf AB, Sweden) at a height of approximately 1.2–1.3 m aboveground. Bore holes in Poland and Spain were plugged with an antifungal mix, while at the remaining sites they were left untreated. Trees for growth analysis were selected following a size-stratified random sampling approach (Jucker, Bouriaud, Avacaritei, Dănilă, et al., 2014), while those for isotope analysis were randomly selected from a subset of dominant and co-dominant trees in each plot (Grossiord et al., 2014) (Fig. 1). These tree cores were used to investigate the effects of forest diversity on productivity and drought resistance (Grossiord et al., 2014; Jucker, Bouriaud, Avacaritei, & Coomes, 2014; Jucker, Bouriaud, Avacaritei, Dănilă, et al., 2014).

For the purposes of this analysis, we only retained data from plots with a complete census record between 2012 and 2022. Moreover, we removed all data from *Q. ilex* trees in Spain, as the majority of these were multi-stemmed and had not been individually tagged in 2012. Consequently, we could not be certain that the same stem was consistently re-surveyed across each census period. We also removed individuals of *Betula pendula* from Finland, as those that were cored were not resurveyed after 2012 due to exhibiting stem damage as a result of the coring itself (see Discussion for more details). In total, this left us with 3201 trees that were cored in 2012 (2416 cored once and 785 cored twice) and 7262 trees that were not cored (Table 1).

### Impacts of tree coring on growth

We assessed the impacts of tree coring on growth using two complementary approaches, the first based on a control–impact design and the second on a before–after comparison (Christie et al., 2019). First, we compared the growth of trees that were cored in 2012 to those that were not cored in the five and ten years after coring took place using the re-survey data from 2017 and 2022. To complement this, for the trees that were cored in 2012 and survived until 2022, we compared their growth rates in the decade before (2001–2011, estimated from tree ring width) and after coring (2012–2022, estimated from measured basal area). In all cases, growth was expressed as a the mean basal area increment (BAI, in cm^2^ year^−1^), as BAI better reflects changes in whole-stem volume growth over the course of a tree’s life compared to radial ring width increments (Klesse & Bigler, 2025). Note that because of measurement errors in DBH that are common with re-census plot data (Talbot et al., 2014), a small subset of trees (7.5%) exhibited negative growth values between 2012 and 2022 (Fig. S1). These negative growth values were mostly concentrated in slow-growing trees from Spain and Italy, suggesting they are most likely the result of random measurement errors that can exceed small stem size increments over these time windows (Fig. S1). Negative growth values were retained in the analysis, as removing them would bias growth estimates as corresponding positive growth errors are much harder to detect and correct (Réjou-Méchain et al., 2017; Talbot et al., 2014).

#### Approach 1: control–impact assessment

To compare the growth rates of trees that were cored to those that were not, we used the census data to fit a hierarchical Bayesian model in which the BAI of tree *i* of species *s* from country *c* in plot *p* was modelled following a skewed normal distribution, which best captured the residual error structure of the model (Fig. S2):

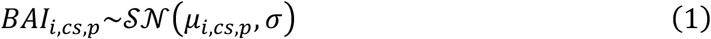

where *μ*_*i*,*cs*,*p*_ is a function of initial tree size (BA_2012_), the crown illumination index (CII) recorded in 2012 (treated as a continuous variable), a categorical variable describing the coring (C) treatment (factor with three levels: not cored, cored once, cored twice), and the interaction term between coring and initial tree size (C × BA_2012_):

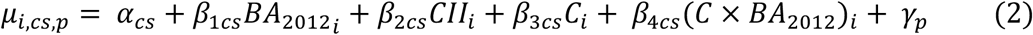

*α_cs_* and *β_1-4cs_* are country- and species-specific model parameters representing the intrinsic growth rate (*α_cs_*), and growth response (i.e., slope) to tree size (*β_1cs_*), CII (*β_2cs_*), coring (*β_3cs_*), and how the effect of coring varies depending on tree size (*β_4cs_*). Additionally, the intercepts were allowed to vary among plots (***γ_p_***) to capture additional variation in BAI among stands. BA_2012_ and CII were included in the model to account for the well-known effects of tree size and competitive status on growth (Jucker *et al*., 2014) allowing us to account for differences in the size distribution of trees that were and were not cored (Fig. 1). The interaction term between coring and tree size was included to test whether smaller trees are disproportionally impacted by coring.

Country-species-specific model parameters (*α_sc_, β_1-4cs_*) were modelled using a multivariate normal distribution:

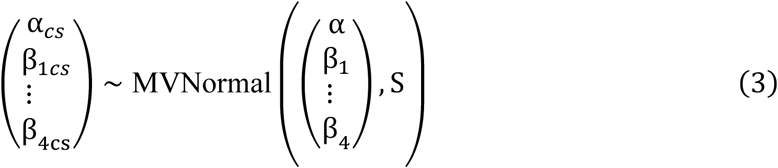

where α represents the community-level intercept, β*_1-4_* the overall effect of covariates on BAI across all species and S is a covariance matrix. Modelling all country-species-level parameters as a multivariate normal distribution allows sharing information across country-species combinations thus improving the fit for poorly represented combinations, while preventing overfitting (McElreath, 2020).

To account for variance in BAI differing among tree species, climatic regions and depending on tree size, we explicitly modelled the residual variance parameter as the following log-linear function (Umlauf & Kneib, 2018):

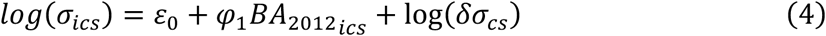

where σ*_isc_* is the residual variance of individual *i* belonging to species *s* from country *c*, ε*_0_* is the intercept, φ_1_ captures how variance changes with tree size, and log(ds*_cs_*) represents the baseline variance for species *s* from country *c*.

#### Approach 2: before–after assessment

To complement the control–impact analysis, we also tested whether trees grew slower after they were cored. To do this, for each tree that had been cored once, we calculated the differences in BAI (ΔBAI) between the decade after coring (BAI_2012-22_) and the one before cores were collected (BAI_2001-11_), where negative values of ΔBAI correspond to slower growth after coring (see Fig. S3 for the same analysis conducted on trees that were cored twice). BAI_2001-11_ was calculated from the annual growth increments measured directly from the cores, whereas BAI_2012-22_ was estimated from the repeat census data. We then modelled ΔBAI of tree *i* belonging to species *s* from country *c* in plot *p* using the same hierarchical Bayesian framework described above but assuming a Student’s t distribution, which best captured the residual error structure of the model (Fig. S4):

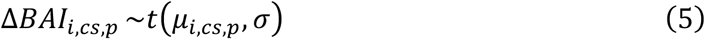

where *μ*_*i*,*sc*,*p*_ is a function of initial tree size (BA_2012_) and the crown illumination index (CII) recorded in 2012 (treated as a continuous variable):

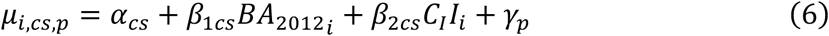

where *α_cs_* and *β_1-2cs_* are country- and species-specific model parameters representing the intercept (*α_cs_*) and ΔBAI response (i.e., slope) to tree size (*β_1cs_*) and crown illumination (*β_2cs_*). Initial tree size and crown illumination were included to account for the potential variation in growth regimes. The intercepts were allowed to vary among plots (***γ_p_*)** to capture additional variation in ΔBAI among stands. Country–species–specific parameters (*⍺*_*sc*_, *β*_1*cs*_, *β*_2*cs*_) were modelled using the same multivariate normal hierarchical structure as described in Eq. (3) for the control-impact assessment growth model.

This analysis makes several assumptions about the comparability of growth trends before and after coring. For example, it overlooks the fact that the climate and the occurrence/severity of extreme events will have differed between the two time periods, thus resulting in differences in growth that have nothing to do with coring. It also ignores the fact that some trees may have experienced growth declines unrelated to coring due to pest and pathogen outbreaks (e.g., spruce bark beetle, ash dieback and the Oriental chestnut gall wasp). Conversely, some trees may have instead experienced increased growth due to competitive release following the mortality of neighbours. Finally, the analysis also assumes that estimates of BAI before and after coring are directly comparable, even though they were obtained using different methodologies (ring width vs repeat DBH measurements). Because radial growth rates can vary noticeably at different positions along the stem, this could potentially introduce considerable noise in the data if tree-cores and DBH measurements were collected at different stem heights, so all data was collected at 1.2-1.3m. To ensure these data were comparable, we also conducted a sensitivity analysis DBH measurements from diameter tapes to those reconstructed directly from the tree ring chronologies shows strong agreement between the two and suggests they directly comparable (Pearson’s correlation coefficient = 0.93; Fig. S5). Despite the limitations of this method, the before–after comparison provides a complementary assessment of the potential impacts of coring on growth.

### Impacts of tree coring on mortality

To assess if coring was associated with an increased probability of mortality, we compared the survival rates of trees that were cored in 2012 to those that were not in the five and ten years after coring took place using the re-survey data from 2017 and 2022. This is analogous to the control–impact assessment on growth described above and made use of the same data and a similar model structure. Specifically, the probability of mortality (*P*_[M=1]_) was expressed as a logistic function of initial tree size (BA_2012_), crown illumination index (CII; continuous variable) and coring treatment (C; factor with three levels: not cored, cored once, cored twice).

*P*_[M=1]_ was modelled as a Bernoulli distribution with a logit link and allowed to vary among species *s*, countries *c* and plots *p* following a hierarchical model structure:

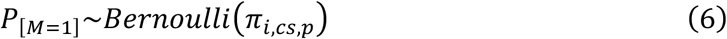

where *π*_*i*,*sc*,*p*_ is a function of initial tree size (BA_2012_) and the crown illumination index (CII) recorded in 2012 (treated as a continuous variable):

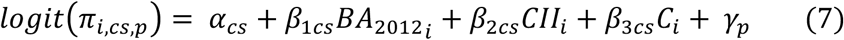

Country–species–specific parameters (*⍺*_*cs*_, *β*_1–3*cs*_) were modelled using the same multivariate normal hierarchical structure as described in Eq. (3) for the control-impact assessment growth model.

An interaction term between BA_2012_ and coring treatment was initially included in the model to test whether coring effects on mortality were most pronounced for small trees. However, it was subsequently removed as we found it was unstable across model runs and did not improve the fit of the model (LOO-adjusted R^2^ = 0.391 with the interaction term and 0.394 without it).

### Model fitting

All models were fit using the *brms* package in R (Bürkner, 2017). For each model, we used four chains and 3000 iterations per chain, with 1500 discarded as warmup. Chains of all models mixed well with R^ = 1.00–1.01. The response variable and all model predictors were scaled and centred prior to model fitting. We used uninformative priors, and model parameter posteriors were summarised through their median and 95% highest-density continuous interval (HDCI) using the *tidyverse* and *tidybayes* packages (Kay, 2024; Wickham et al., 2019). Predictors were considered to have a clear effect on the response variable when the HDCI of their model coefficients did not encompass zero.

Model predictions plotted in Figs 2 and 4 were created using the *posterior_linpred()* function in the *brms* R package (Bürkner, 2017). This method describes the uncertainty at the group level and does not include individual level variation (σ).

**Fig. 2:**
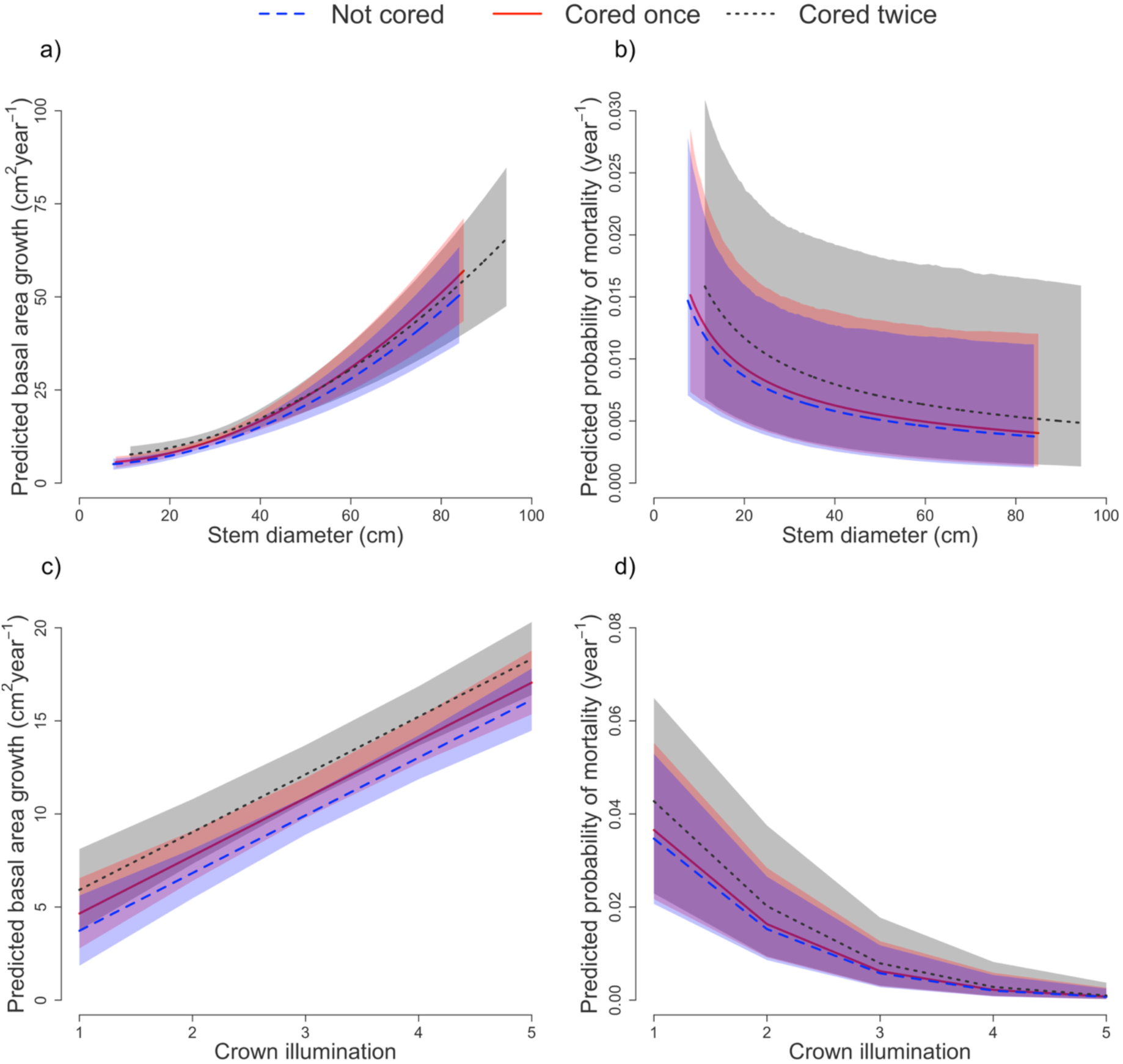
Differences in basal area growth (left) and mortality rates (right) for trees that were either not cored (blue, dashed line), cored once (red, continuous line) or cored twice (black, dotted line) across a range of stem diameter (**a**–**b**) and crown illumination values (**c**–**d**). Fitted lines are model predictions with 95% highest posterior density intervals. For panels (**a**–**b**) a crown illumination value of three was used to generate the predictions, while for (**c**–**d**) stem diameter was fixed at 25.7 cm (quadratic mean value across the entire dataset).

## Results

### Impacts of tree coring on growth

#### Approach 1: control–impact assessment

Variation in BAI among trees was strongly and positively associated with both tree size (Fig. 2a) and crown illumination index (Fig. 2c), with larger, canopy-dominant trees exhibiting noticeably faster growth rates. On average, a tree with a DBH of 60 cm had a mean BAI that was 4.2 times greater than that of a tree with a DBH of 15 cm (29.9 cm^2^ year^−1^ vs 7.1 cm^2^ year^−1^, assuming CII = 3 in both cases).

When we compared the BAI of trees that had been cored with those that were not, we found only minimal differences in growth rate and no evidence to suggest that coring detrimentally affected BAI in either the five or ten years after coring (Fig. 2 and Figs S6–7). The predicted growth for a tree of average size (DBH = 25.7 cm, quadratic mean diameter across all surveyed trees) that was not cored was 10.0 cm^2^ year^−1^, slightly lower than that of a tree that was cored once (11.0 cm^2^ year^−1^; assuming CII = 3 in both cases). This positive growth response to coring was slightly more pronounced for trees that were cored twice (BAI = 11.2 cm^2^ year^−1^).

We also found no evidence to suggest that coring adversely impacted the growth of smaller trees more than others, with no clear interaction term between coring and tree size in the models (Figs S6–7). Moreover, the effects of coring on growth were remarkably consistent across tree species and forest types (Fig. 3a), including when we compared broadleaves and conifers.

**Fig. 3:**
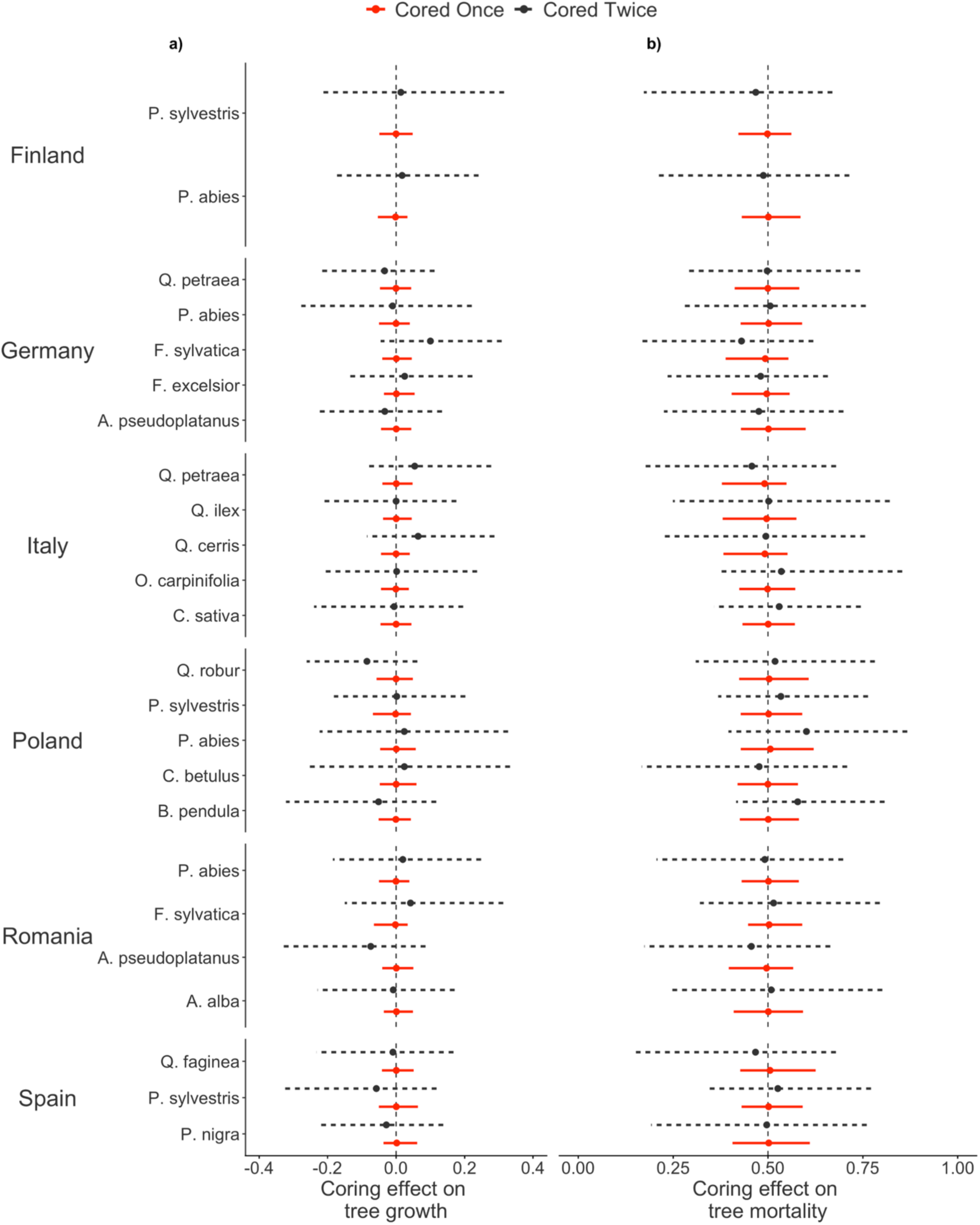
Variation in the effects of tree coring on **(a)** growth and **(b)** mortality rates across species and study sites for trees that were cored once (red, continuous line) or twice (black, dotted line), expressed as the difference relative to trees that were not cored. For growth, points represent posterior means of the standardised model coefficients (±95% highest posterior density intervals), while for mortality they show the probability of a cored tree dying compared to non-cored tree (where 0.5 represents no difference between the two).

#### Approach 2: before–after assessment

When we compared the BAI of cored trees in the decade before and after coring took place in 2012, we again found no indication of an adverse growth response to coring (Fig. 4). Tree growth rates in the decade before and after coring were strongly positively correlated (Pearson’s correlation coefficient = 0.70; Fig. 4a). Consistent with our comparison of cored and non-cored trees, we observed that trees exhibited slightly elevated BAI in the decade after coring (ΔBAI = 0.14 cm^2^ year^−1^ for a tree with DBH = 25. 7 cm and CII = 3). Once again, these results were largely consistent across species and forest types (Fig. 5). We did however observe that in certain sites, ΔBAI was systematically greater (Poland) or lower (Spain) in the decade following coring.

**Fig. 4:**
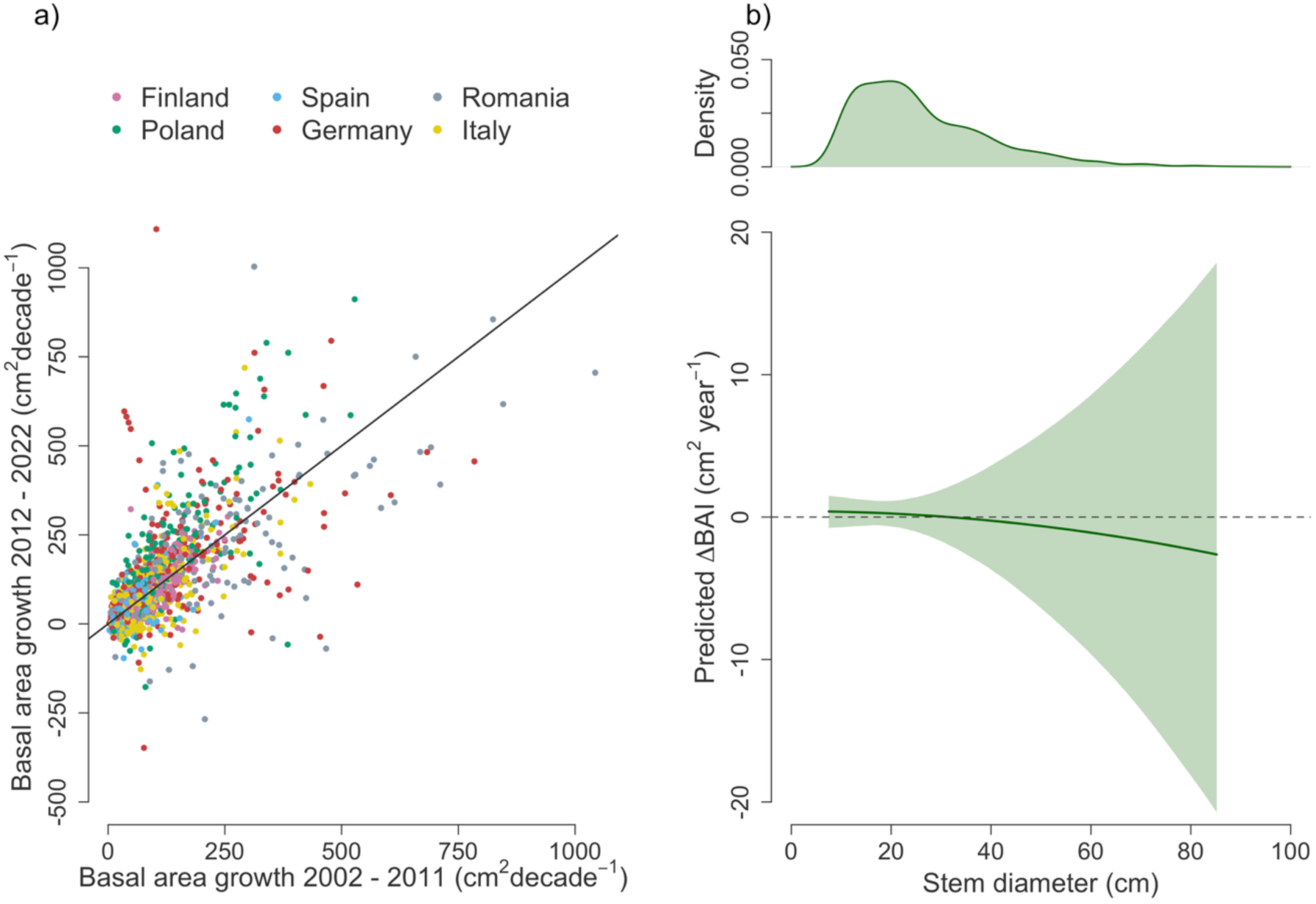
Tree growth rates before and after coring. Panel (**a**) shows the relationship between the basal area growth of trees cored once in 2012 in the decade before and after coring (calculated from the tree ring and census data, respectively). Points are coloured by country, and the line corresponds to a 1:1 relationship. Panel (**b**) shows the estimated difference (95% highest posterior density intervals) in basal area increment before and after coring (ΔBAI) as a function of tree size, where values of 0 correspond to no difference in growth over time. The density plot at the top illustrates the size distribution of trees included in the analysis.

**Fig. 5:**
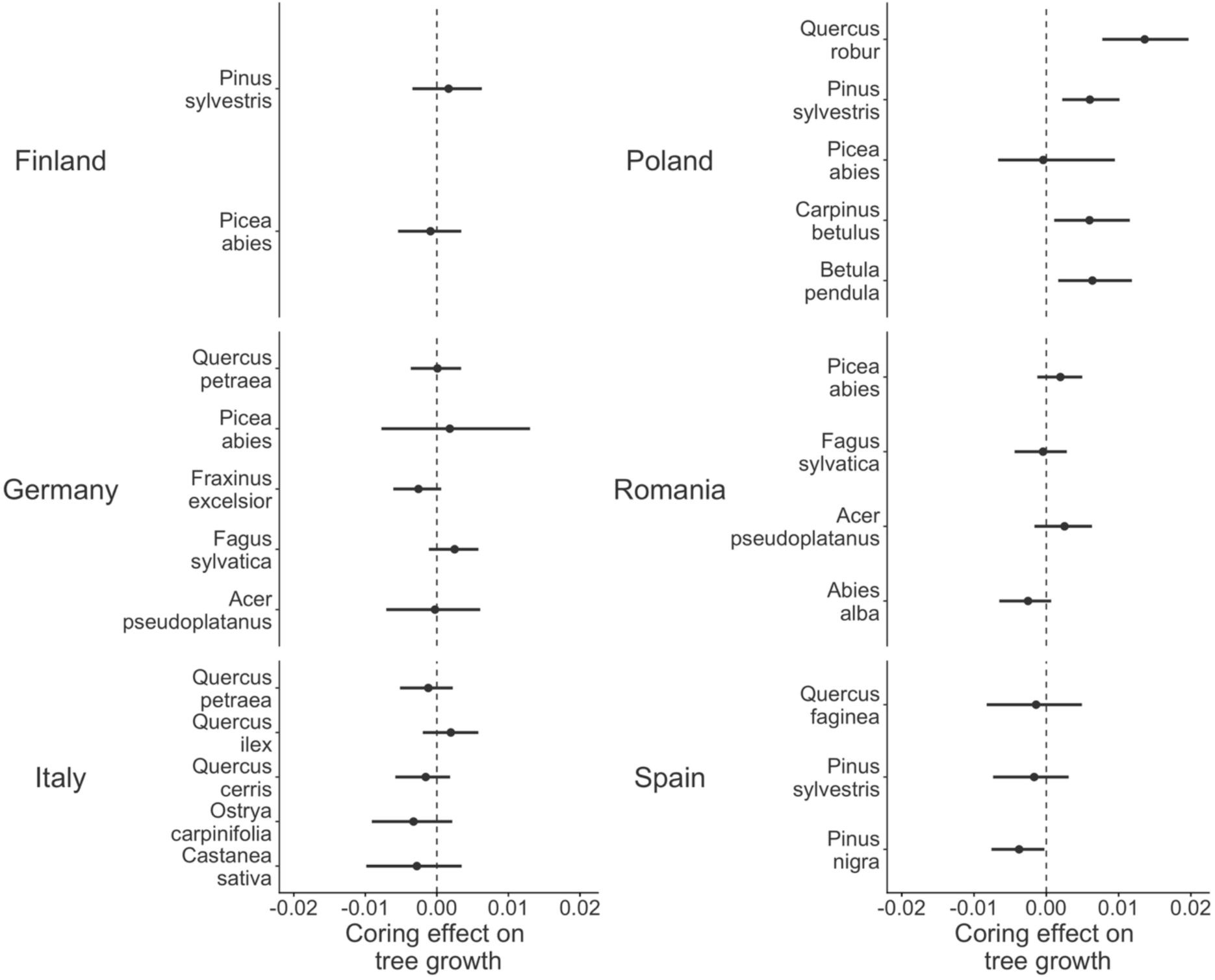
Variation growth rates before and after coring across species and study sites for trees that were cored once. Points are standardised model coefficients with 95% highest posterior density intervals. A negative value would indicate that cored trees grew slower in the decade after coring.

### Impacts of tree coring on mortality

Regardless of whether they were cored, 20.0% of trees across the 2012–2022 census period died (Table 1), with the highest mortality rates occurring in Germany and Italy (24.8% and 24.6% across species, respectively), with *Picea abies* in Poland and Germany having the highest species-level mortality rate (68.2% and 66.9% respectively). Similarly to growth, mortality rates varied with both initial tree size (Fig. 2b) and crown illumination index (Fig. 2d), with larger, canopy-dominant trees exhibiting the lowest mortality rates. On average, a tree with a DBH of 60 cm and a CII of 5 (large, canopy dominant tree) had a probability of dying of 0.07% year^−1^ compared to 4.7% year^−1^ for a tree with a DBH of 15 cm and a CII of 1 (small-sized understory tree).

Cored trees had a slightly elevated mortality risk in the decade after coring (Fig. 2 and Fig. S6), although the uncertainty around these estimates was very large and fully overlapped with zero. Specifically, a tree of average size (DBH = 25.7 cm, CII = 3) that had not been cored had an expected probability of mortality of 0.75% year^−1^ (95% highest posterior density intervals = 0.38–1.45%), which increased to 0.81% year^−1^ (0.41–1.56%) for a tree that was cored once and 1.02 % year^−1^ (0.42–2.16 %) for trees cored twice. These results were almost identical when only considering trees that died in the five year period after coring, up to 2017 (Fig. S7). They were also largely consistent across forest types and tree species (particularly for trees cored once, for which sample sizes were larger and uncertainly lower; Fig. 3b), with no clear evidence to suggest that certain functional groups were more prone to dying after coring than others.

## Discussion

Using plot census and tree ring data from the FunDivEUROPE network, we quantified the effect of increment coring on tree growth and mortality across Europe’s dominant tree species. This study provides the most comprehensive assessment to date on the impacts of coring across a broad range species and forest types, including Europe’s Mediterranean, temperate and boreal forests. We found coring had no negative impact on tree growth, while mortality responses were much more uncertain. Our data enabled us to test whether responses to coring vary with tree size and evaluate whether standard dendrochronological practices, such as extracting multiple cores per tree, are associated with long-term impacts on tree growth and mortality.

### No evidence of adverse effects of tree coring on growth

Our results indicate that coring had no negative effect on tree growth in the decade after cores were extracted. This is consistent with the small number of previous studies that have characterised growth responses to coring (Fabiánová & Šilhán, 2021; Mantgem & Stephenson, 2004; Portier et al., 2023; Wunder et al., 2011), but significantly build on this evidence by expanding the scope to include many more species and forest types. While we observed no declines in growth following coring, we did detect a positive growth response which was most pronounced in twice-cored trees and exceed typical measurement error associated with diameter tapes (Luoma et al., 2017). This raises the question of whether these patterns reflect a biological response to coring or if instead they are a result of sampling bias toward coring larger, canopy dominant trees that were not fully accounted for in our statistical models.

Localised positive growth responses have been reported previously for *P. abies* as a result of vertical scarring (Fabiánová & Šilhán, 2021), and coring has also been shown to induce pronounced wound responses in some species (Portier et al., 2023). As trees in our study were cored at a similar height at which stem diameters are measured, increased growth may therefore result from a wound response to the coring, with a second wound amplifying this effect. Alternatively, the response may be driven by hormesis, whereby a low dose of stress elicits a stimulatory response (Agathokleous et al., 2019). Although hormesis is reported to occur widely in plants subjected to physical stressors (Agathokleous et al., 2019, 2020; Salinitro et al., 2021), evidence in mature trees is currently lacking and would need further exploration in the context of coring. Irrespective of the underlying mechanism, any growth response to coring is likely to be localised around the coring site (Fabiánová & Šilhán, 2021; Tsen et al., 2016). If not accounted for, these localised responses to coring risks inflating diameter measurements, potentially leading to overestimates of growth and biomass in permanent monitoring plots. We therefore recommend that increment cores be extracted at least 30 cm above or below the point at which diameter measurement are taken as a precautionary measure.

We found no evidence that small trees were any more vulnerable to coring than large ones in terms of growth, regardless of whether they were cored once or twice. Coring wounds are relatively small compared to the minimum size of the stem in this study (10 cm diameter for cored trees), and trees have a high resilience to the loss of xylem (Dietrich et al., 2018). While the immediate hydraulic impacts of coring are therefore likely negligible, we cannot rule out potential longer-term consequences for smaller trees as internal scarring from coring may predispose stems to internal decay (Tsen et al., 2016), which can disproportionately impact smaller individuals (Datta et al., 2025). Therefore, minimum diameter thresholds for coring should be considered, accompanied by longer-term monitoring of wound sites for evidence of decay.

Growth responses to coring were also very similar across tree species and forest types. Previous work has suggested that gymnosperms may be more resilient to coring than angiosperms (Tsen et al., 2016) this may be due to their ability to close wounds with sap and resin, but we found no clear evidence to support these differences among functional groups. A common concern with coring is that wounds may act as a vector for pathogens (Tsen et al., 2016), particularly fungi that cause wood decay. If so, we would have expected to see stronger responses to coring in regions with milder and wetter climates, but again this did not emerge from the data. In this regard it is useful to keep in mind that coring holes are much smaller than the typical wound caused by the loss of a branch, something which trees will experience frequently across their lifetimes. This suggests that coring is unlikely to significantly increase a tree’s exposure to pathogens, irrespective of the climate or species. Nevertheless, to better resolve this question would warrant specifically monitoring coring wounds for evidence of infection and wood rot.

Our analysis of the impacts of tree coring on growth comes with some important limitations. Studies explicitly designed to test the effects of coring on trees are rare, and like most others ours was performed retrospectively and opportunistically. Post-hoc approaches make testing for the impact of coring a non-trivial task as selection bias and unmeasured confounding factors cannot be fully accounted for. These challenges are particularly evident in the before-after growth comparisons, which is complicated by differences in methodology of measuring growth (cores vs repeated stem diameter measurements) and by the challenge of comparing growth across time periods with contrasting climates. Comparing cored and uncored trees also present its challenges, particularly when it comes to ruling out potential selection biases towards coring larger and potentially healthier trees. While our statistical models explicitly account for differences in tree size and crown illumination, other factors that relate to tree vigour may not have been captured. As a result, the positive growth responses associated with coring may partially reflect the preferential sampling (conscious or unconscious) of healthier individuals for coring rather than a biological response to coring per se.

### Effects of tree coring on mortality remain uncertain and require further investigation

In comparison to growth responses to coring, effects on mortality were much more uncertain. Model estimates indicate that on average cored trees had slightly lower survival rates that their uncored counterparts, an effect that was more pronounced for trees cored twice. However, these estimates came with a large degree of uncertainty, with posterior density intervals overlapping widely across coring treatments. This underscores the general challenge of attributing what factors modulate tree mortality and by how much (Hammond et al., 2022; International Tree Mortality Network, 2025). This is an inevitable consequence of attempting to detect statistical signals for processes that are inherently rare and stochastic such as mortality, which on average only affects 1-2% of trees per year and is often characterised by punctuated distributions through time that require very large and temporally resolved datasets to detect with certainty (Araujo et al., 2021; Lines et al., 2010; Zuleta et al., 2022).

Complicating things further, in certain species possible increases in mortality risk due to coring could have been masked due to baseline mortality rates being elevated by other factors, such as bark beetle outbreaks in *Picea abies*, ash dieback in *Fraxinus excelsior* in Germany and the oriental chestnut gall wasp infesting *Castanea sativa* in Italy (Fuchs et al., 2024; Lausch et al., 2013; Seidl et al., 2014).

While our data indicate that substantial increases in mortality due to coring are very unlikely, we cannot fully rule out more subtle influences that would require larger sample sizes to be detected with confidence. Even assuming that mortality rates are slightly elevated from coring, given that only a relatively small subset of trees are typically selected for coring and because background mortality rates are generally low, in practise only a very small number of additional trees would potentially die due to coring. Based on the estimates from this study, having cored ∼30% of around 10,500 trees – an unusually high proportion – would have resulted in approximately 1-2 additional tree deaths per year across the entire network. Nevertheless, decisions on whether to core specific trees should take this risk into account, particularly if coring individuals of conservation and/or cultural value and if choosing to extract multiple cores per trees.

More specifically, it is important to acknowledge that there are circumstances under which tree coring can have deleterious outcomes for trees. For example, as mentioned in the Methods section, several birch trees (*Betula pendula*) that were cored in Finland are known to have subsequently suffered stem cracking and canopy dieback in the years prior to the 2017 plot census. This was severe enough that a decision was made to stop monitoring these individuals in 2017 and establish new plots to replace them, which is why they were not included in our analysis. Based on our own personal observations in this field, the stem cracking appears to have resulted as a consequence of freeze thaw cycles over winter following water becoming trapped in the bore hole. Bore holes were noted to have a slight downward inclination, facilitating water pooling. Moreover, coring took place late in the growing season (mid-September), giving trees little opportunity to heal wounds before the onset of winter – which occurs early and is prolonged in the boreal zone. It is unlikely that this adverse effect of coring reflects a general predisposition of birch to coring, as we did not observe the same response for birch trees cored in Poland earlier that same summer. This highlights how context specific responses to coring can be, making them inherently harder to predict.

### Recommendations for future work

Despite our study being among the most comprehensive to date to test the impacts of coring on tree growth and survival, our results point to several important avenues for future research needed to inform the use of tree coring in ecological research. In terms of expanding sample sizes to better address uncertainty around how coring influences a tree’s long-term mortality risk, one possible avenue would be to leverage large-scale national forest inventories that integrate coring as part of their monitoring programs, such as those of Canada, France and Sweeden among others. Similarly, to help better understand the potential role of climate in modulating the risk of wood decay following coring, studies focusing on the growing use of dendrochronology in the tropics are warranted. The handful of studies that have explored the impacts of coring in tropical regions have so far reported no negative outcomes (Neo et al., 2017; Palakit & Pumijumnong, 2024), but expanding these to cover a broader range of species and environmental contexts remains a priority.

In addition to this opportunistic approach to studying the effects of coring after the fact, there remains a clear need to design targeted field experiments that aim to unpack some of the important nuance around tree responses to coring. These studies should follow robust Before-After Control-Impact (BACI) designs by leveraging paired cored and non-cored trees. This design would minimise confounding effects and could be used to further probe important questions around the efficacy of specific wound treatment protocols, the influence of bore hole size (e.g., 5 vs 10 mm diameter) on risk of infection and stem damage, and how timing of coring during the tree’s growth cycle influences outcomes. Critically, a key feature of these targeted studies should be the implementation of frequent and long-term monitoring of trees after coring to assess wounds for scarring and signs of decay (Wunder et al., 2013), as well as building continuous mortality curves to identify when trees actually die (Zuleta et al., 2022). Monitoring protocols should also aim to go beyond relatively crude measures of tree health based on growth and survival, incorporating other key aspects of tree vitality such as crown condition and dieback. Together, these approaches would bring further clarity around the potential impacts of coring on trees, allowing this important and versatile research tool to be used safely and with confidence across a range of disciplines.

## Acknowledgements

The FunDivEUROPE project was funded through the European Union Seventh Framework Programme (grant: 265171). This work was funded by a UKRI Frontier Research award (grant: EP/Y003810/1) awarded to TJ, which also supported RB and DN.

## Author contribution statement

OB and GI coordinated the establishment and re-census of the FunDivEUROPE plot network. TJ generated the tree ring data, with the assistance of OB. RB and TJ designed the study with input from TSO. RB analysed the data, with assistance from DN and FJF. RB wrote the first draft of the manuscript with the assistance of TJ and TSO. All authors contributed substantially to revisions.

## Data and code availability statement

All data and R code underpinning the results of this study will be publicly archived on Zenodo following the review of this paper.

