## Supplementary material for "Coring has no detrimental impact on tree growth across European forests, but effects on mortality remain uncertain"

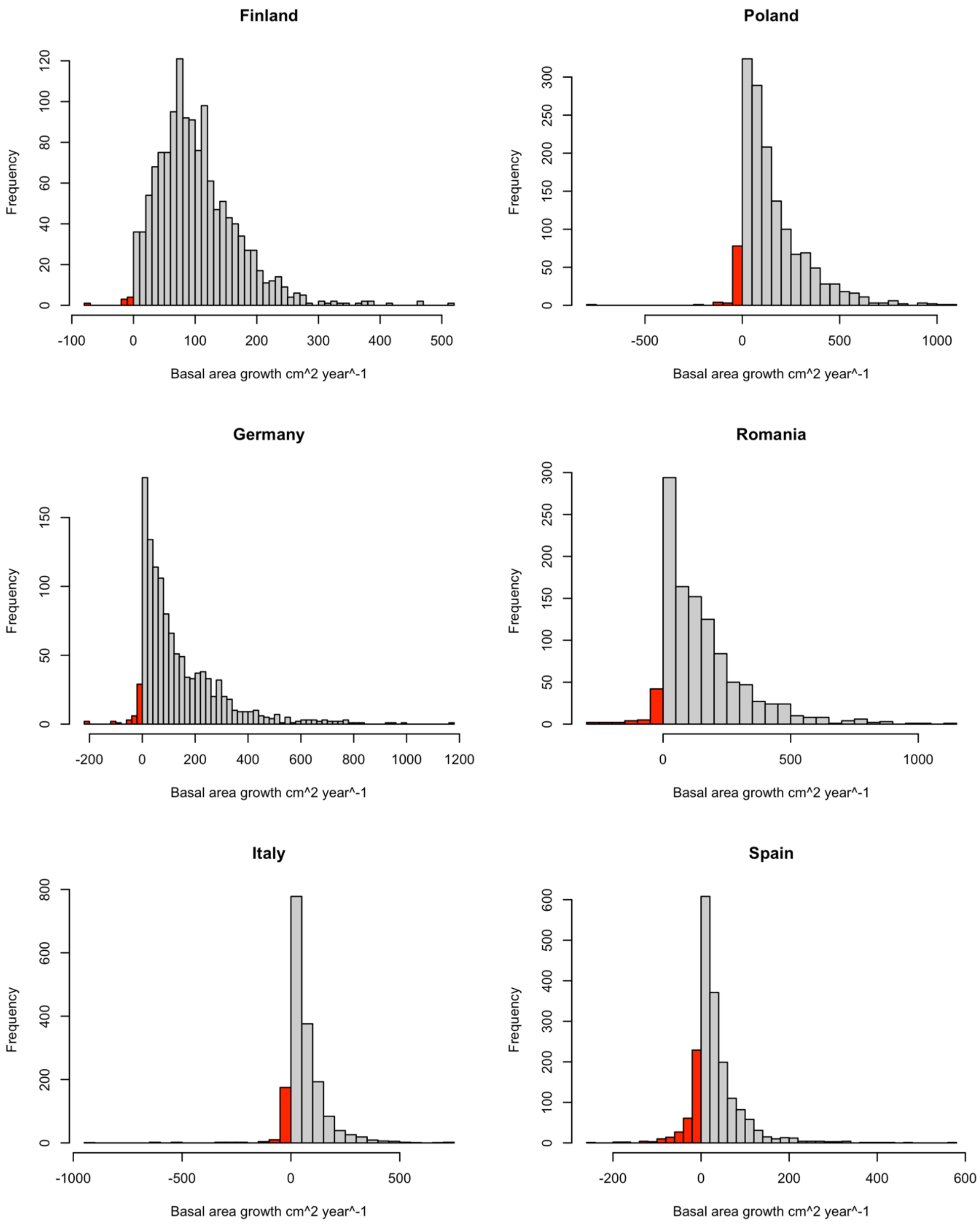


**Fig S1**: Distribution of growth values by country, with negative growth values shown in red.

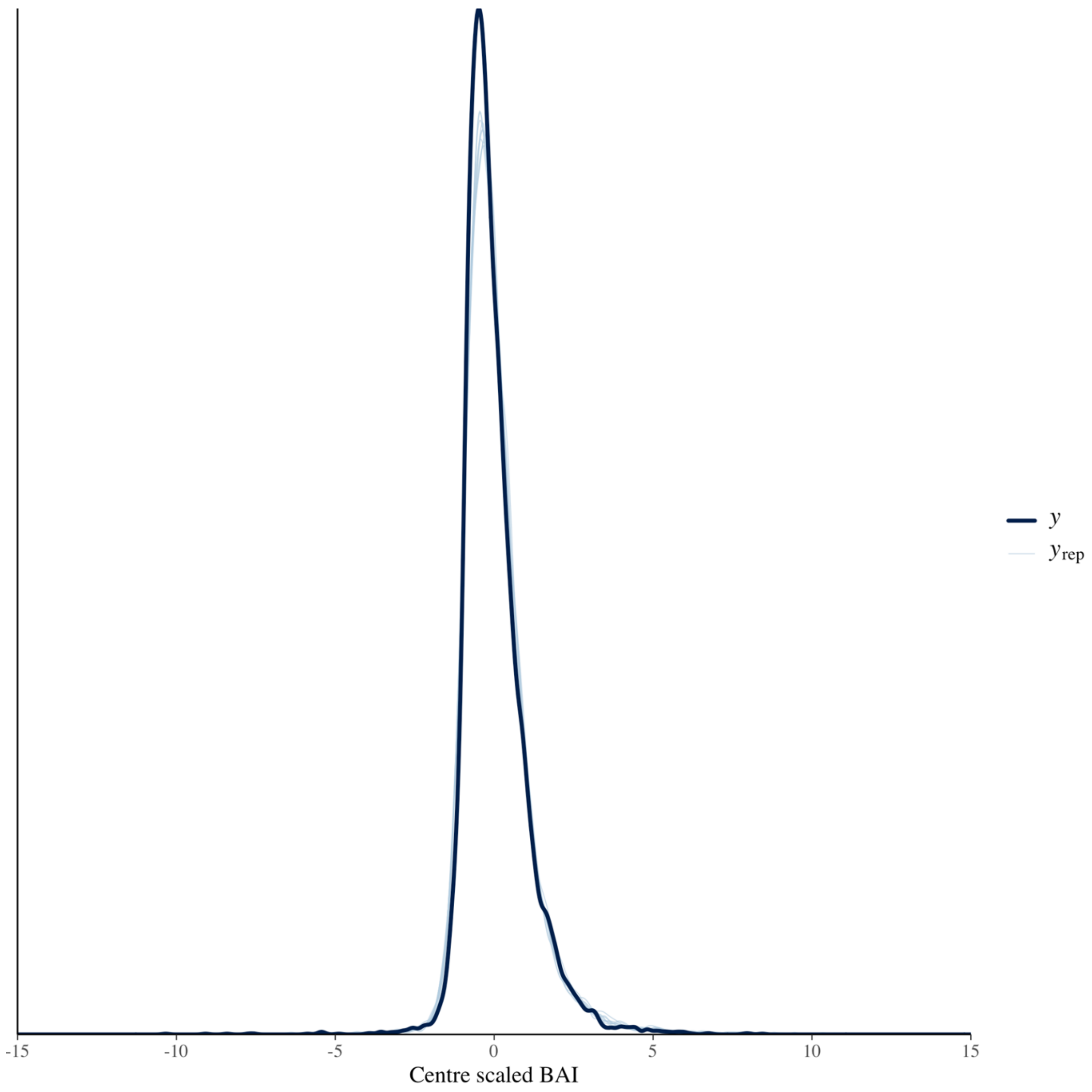


**Fig S2:** Comparison of the observed data to the posterior predicted distribution for the 10 year control-impact growth model, with a skewed normal error distribution. The black line is the observed data while the blue lines are simulated from the posterior distribution.

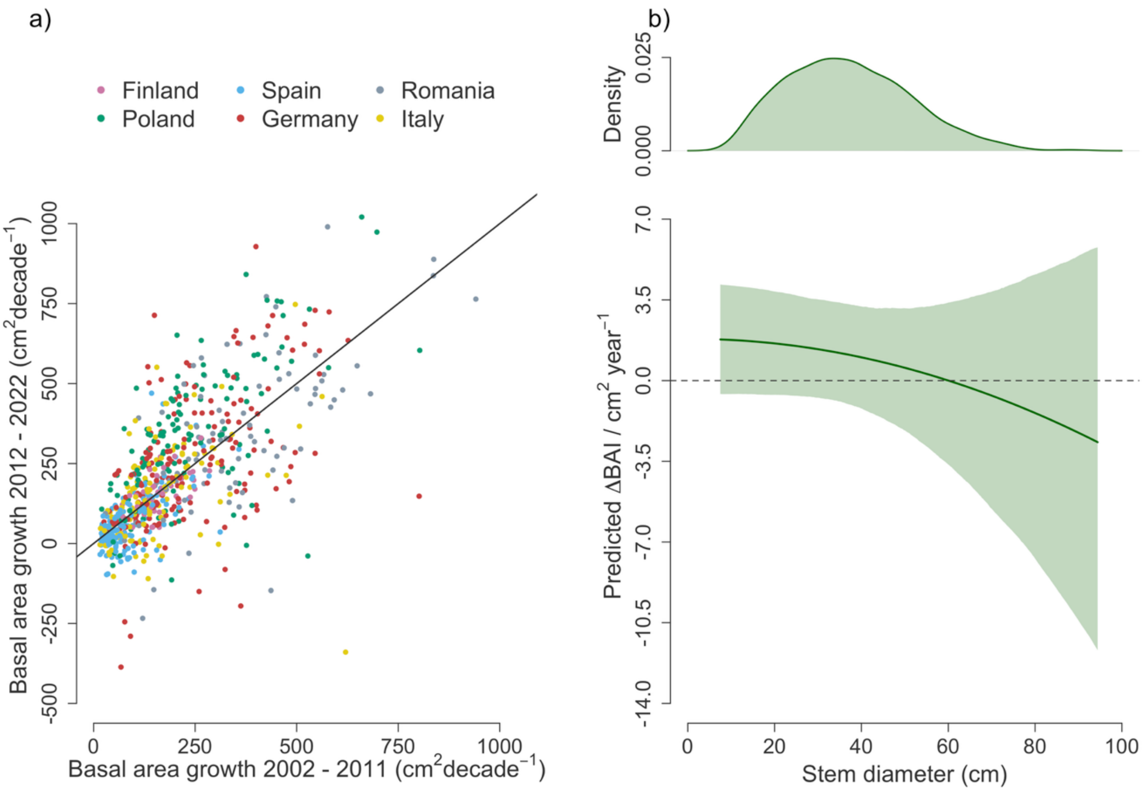


**Fig S3**: Tree growth rates before and after coring for trees that were cored twice. Panel (a) shows the relationship between the basal area growth of trees cored twice in 2012 in the decade before and after coring (calculated from the tree ring and census data, respectively). Points are coloured by country and the line corresponds to a 1:1 relationship. Panel (b) shows the estimated difference (95% highest posterior density intervals) in basal area increment before and after coring (ΔBAI) as a function of tree size, where values of 0 correspond to no difference in growth over time. The density plot at the top illustrates the size distribution of trees included in the analysis.

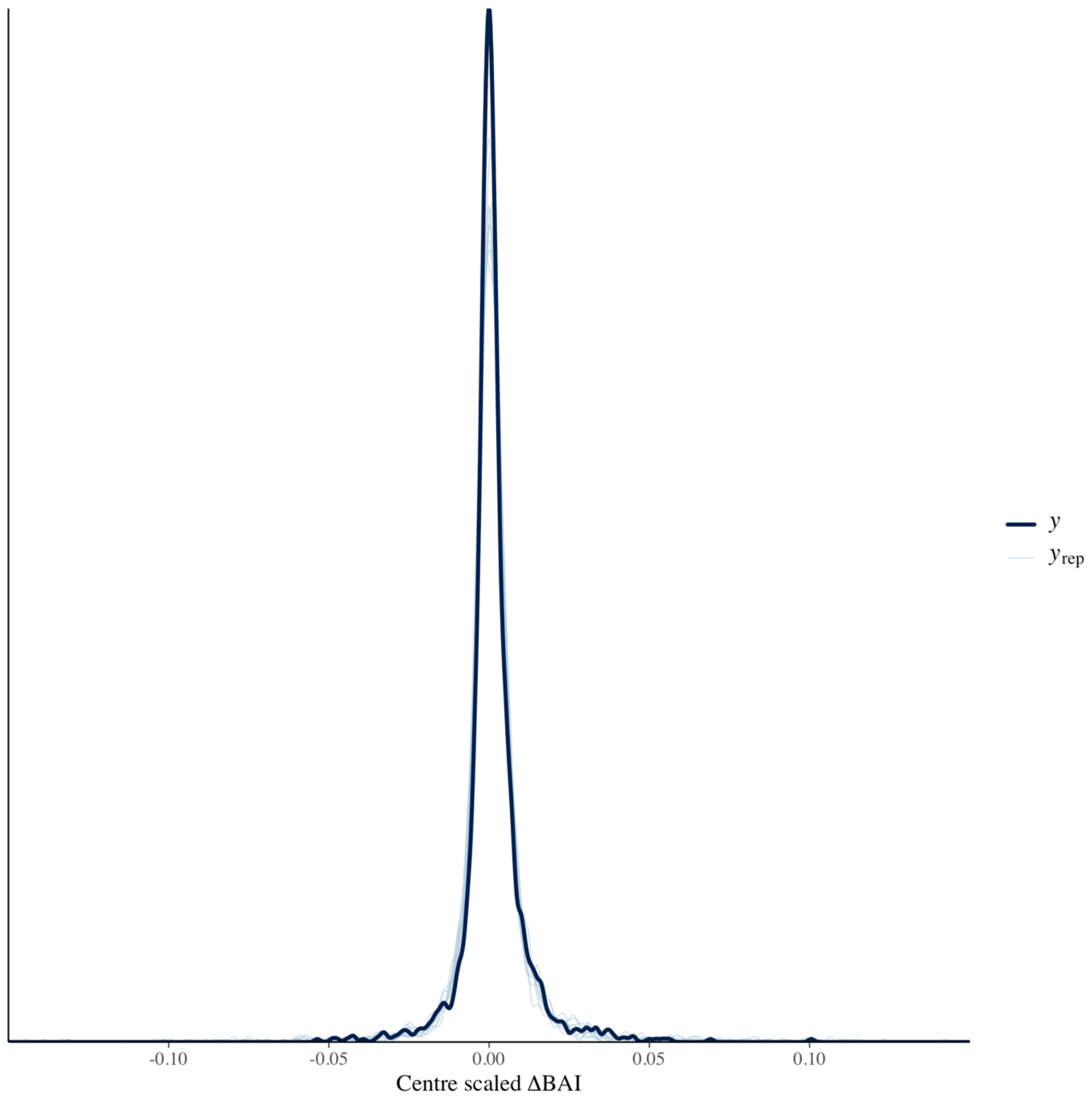


**Fig S4:** Comparison of the observed data to the posterior predicted distribution for the before-after ΔBAI model, with a Student's t error distribution. The black line is the observed data while the blue lines are simulated from the posterior distribution.

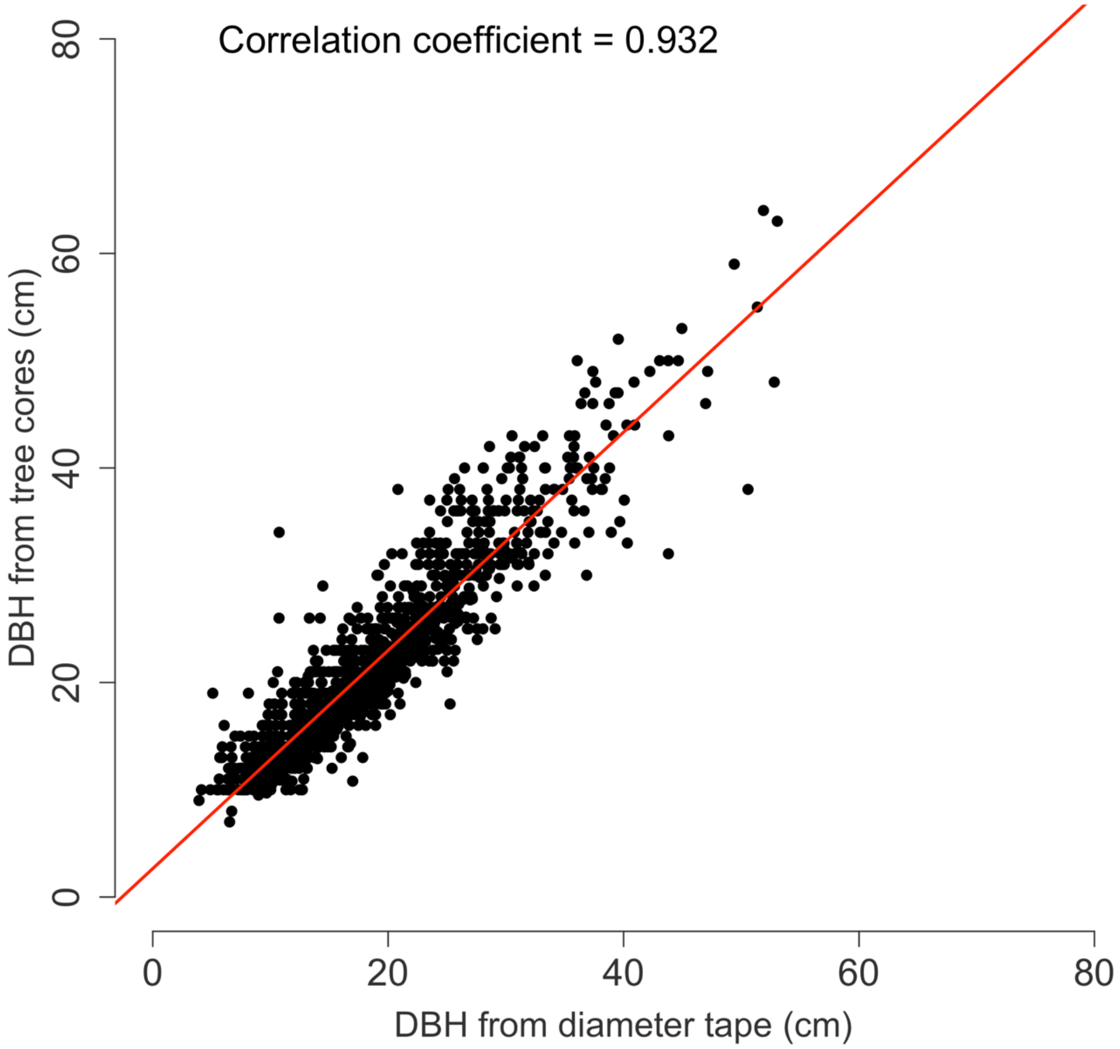


**Fig. S5:** Comparison of DBH measurements obtained from diameter tapes and those reconstructed by sequqentially summing the ring widths of cored trees across their entire lifetime. For this we only used the 1302 trees for which we had complete cores for which distance to pith from the final measured ring was measured. The Pearson correlation coefficient between the two is reported in top right.

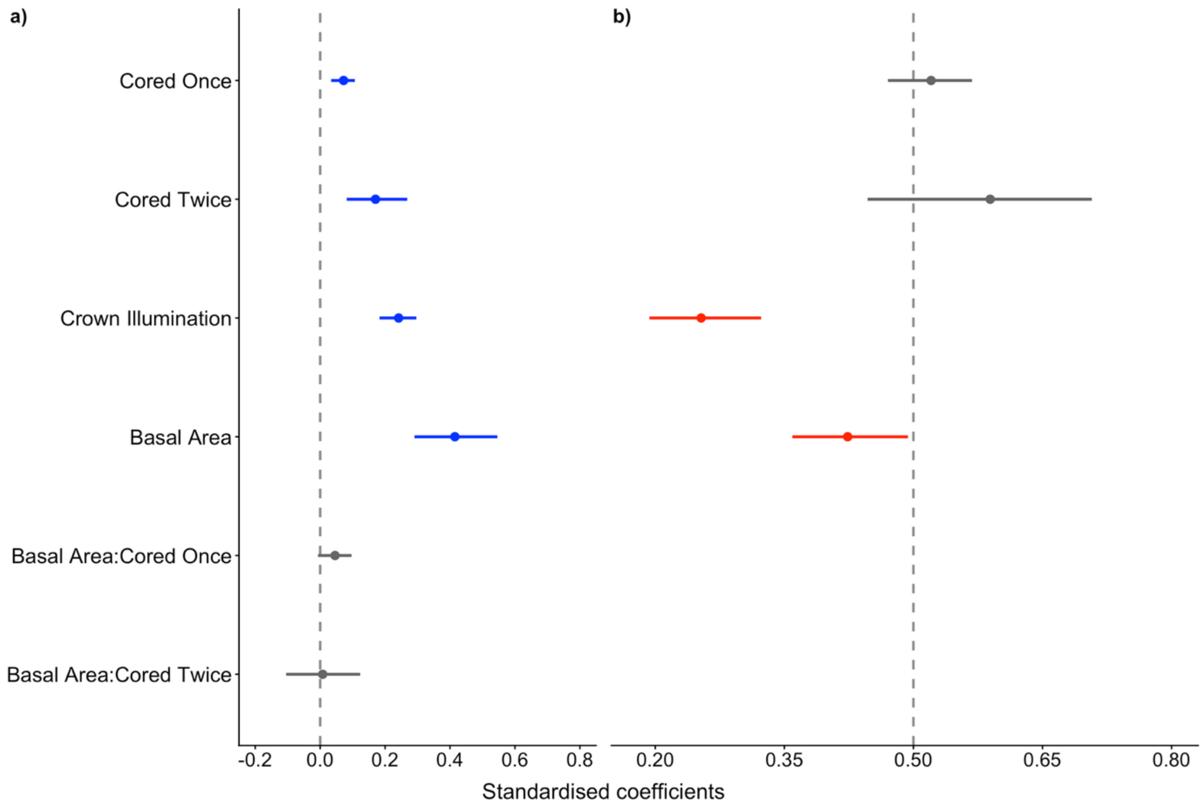


**Fig S6:** Standardized model coefficients for (**a**) tree growth and (**b**) probability of mortality across the 10 year census interval (2012–2022). Standardised model coefficients are shown for growth from the control-impact assessment. For mortality the probability of a cored tree dying compared to non-cored tree is shown, where 0.5 represents no elevated mortality as a result of coring. Circles show the posterior means and lines represent the 95% highest posterior density intervals.

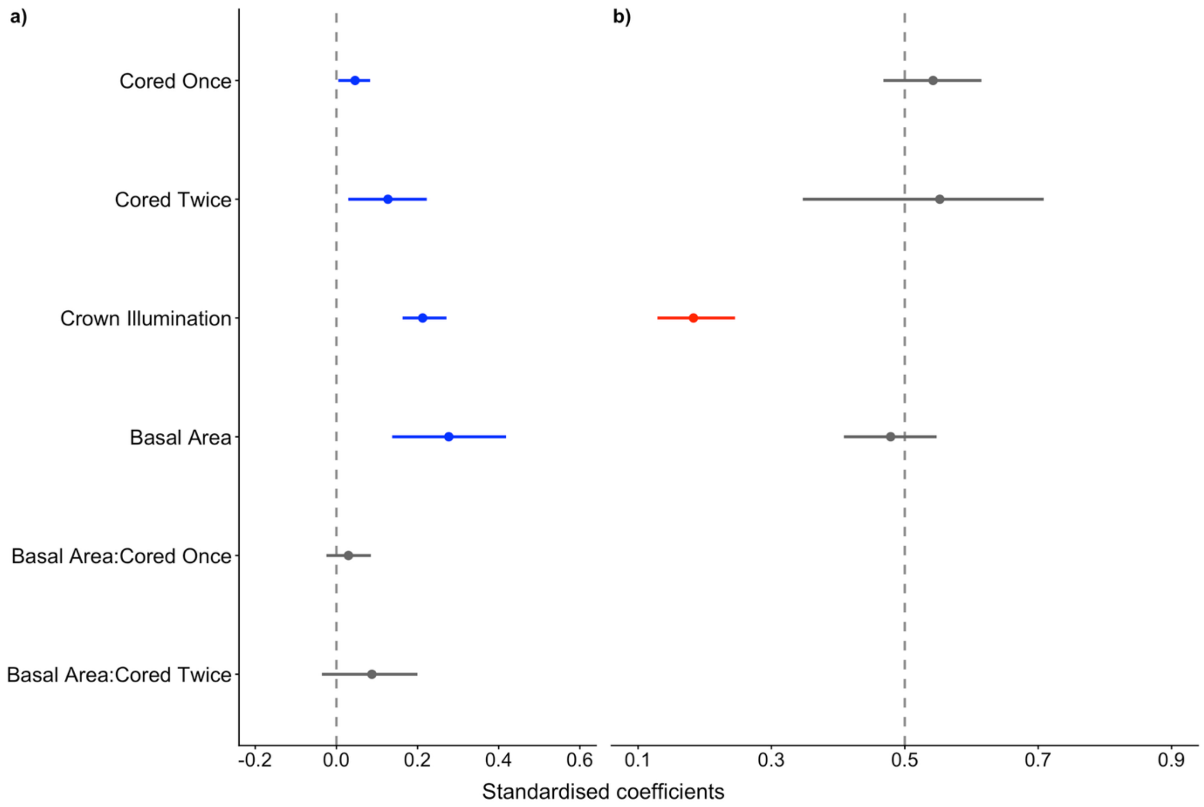


**Fig S7:** Standardized model coefficients for (**a**) tree growth and (**b**) probability of mortality across the 5 year census interval (2012–2017). Standardised model coefficients are shown for growth from the control-impact assessment. For mortality the probability of a cored tree dying compared to non-cored tree is shown, where 0.5 represents no elevated mortality as a result of coring. Circles show the posterior means and lines represent the 95% highest posterior density intervals.

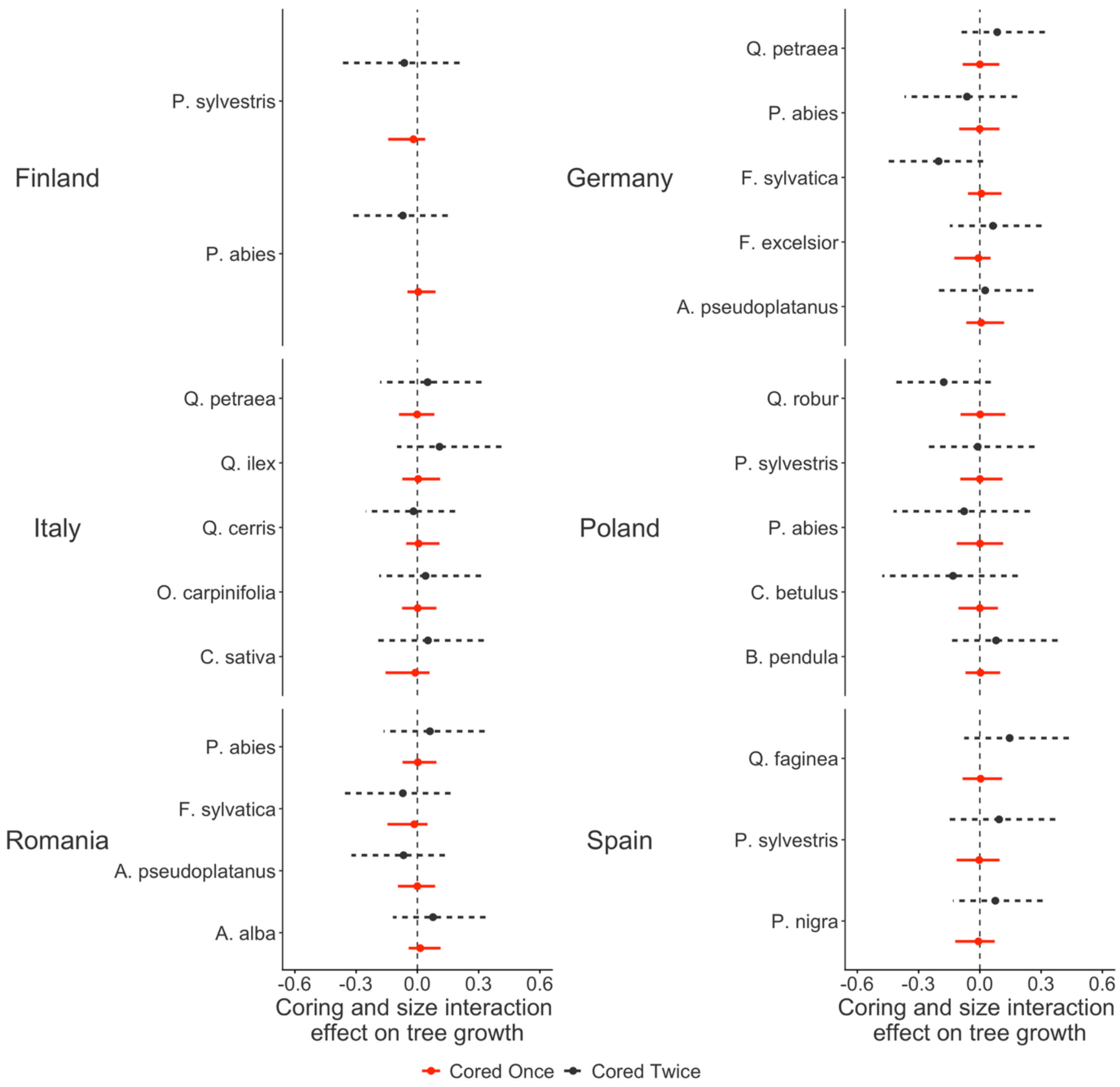


**Fig. S8**: Variation in the effects of the interaction between coring and tree size on growth across species and study sites for trees that were cored once (red, continuous line) or twice (black, dotted line), expressed as the difference relative to trees that were not cored. Points represent posterior means of the standardised model coefficients (95% highest posterior density intervals).

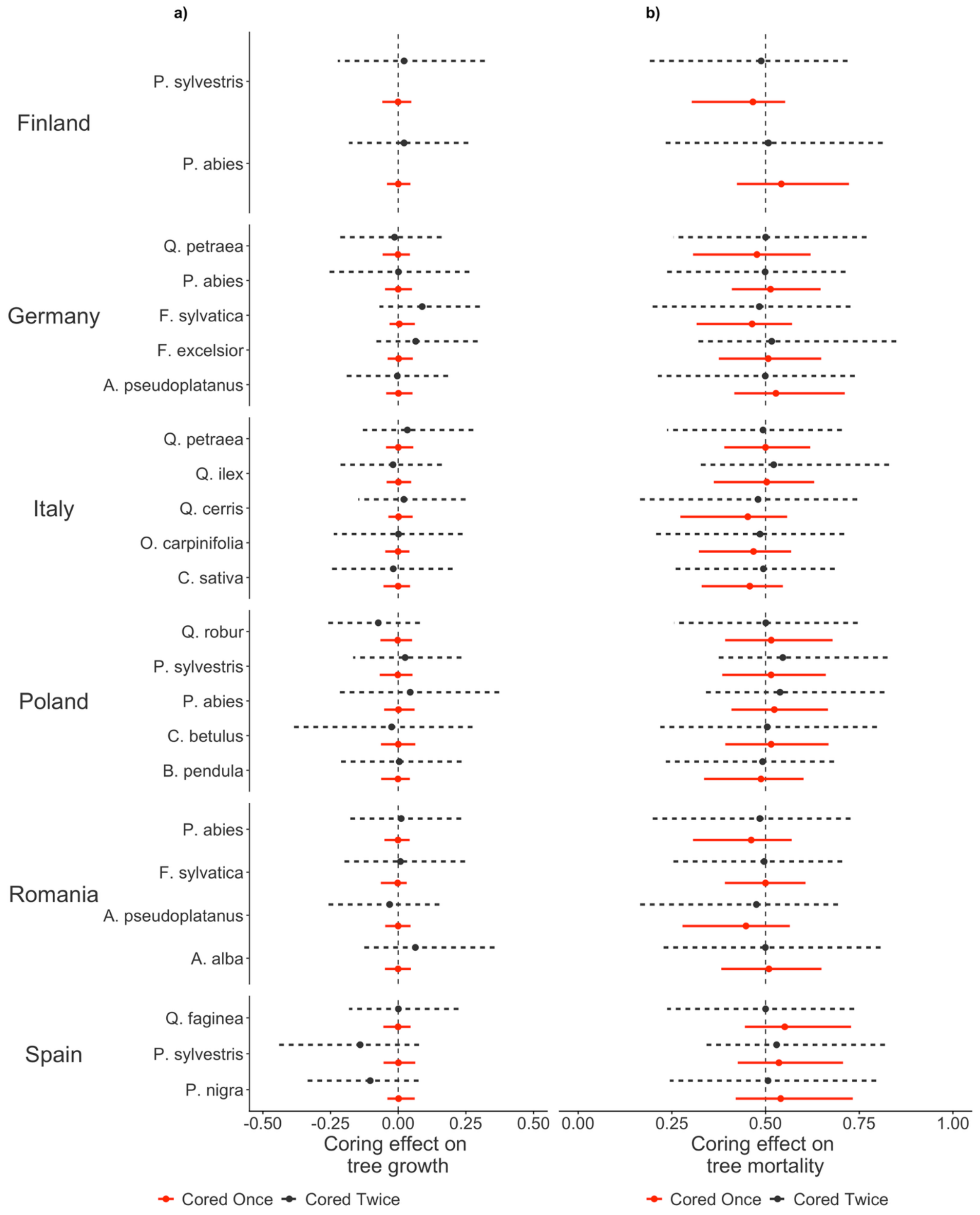


**Fig. S9**: Variation in the effects of tree coring over the 5 year census period (2012-2017) on **(a)** growth and **(b)** mortality rates across species and study sites for trees that were cored once (red, continuous line) or twice (black, dotted line), expressed as the difference relative to trees that were not cored. For growth, points represent posterior means of the standardised model coefficients (95% highest posterior density intervals), while for mortality they show the probability of a cored tree dying compared to non-cored tree (where 0.5 represents no difference between the two).
